# Comparison of breeding strategies for the creation of a synthetic pig line

**DOI:** 10.1101/2021.09.22.461330

**Authors:** Audrey Ganteil, Torsten Pook, Silvia T. Rodriguez-Ramilo, Bruno Ligonesche, Catherine Larzul

## Abstract

Creating a new synthetic line by crossbreeding means complementary traits from pure breeds can be combined in the new population. Although diversity is generated during the crossbreeding stage, in this study, we analyze diversity management before selection starts. Using genomic and phenotypic data from animals belonging to the first generation (G0) of a new line, different simulations were run to evaluate diversity management during the first generations of a new line and to test the effects of starting selection at two alternative times, G3 and G4. Genetic diversity was characterized by allele frequency, inbreeding coefficients based on genomic and pedigree data, and expected heterozygosity. Breeding values were extracted at each generation to evaluate differences in starting selection at G3 or G4. All simulations were run for ten generations. A scenario with genomic data to manage diversity during the first generations of a new line was compared with a random and a selection scenario. As expected, loss of diversity was higher in the selection scenario, while the scenario with diversity control preserved diversity. We also combined the diversity management strategy with different selection scenarios involving different degrees of diversity control. Our simulation results show that a diversity management strategy combining genomic data with selection starting at G4 and a moderate degree of diversity control generates genetic progress and preserves diversity.

## 1 Introduction

Genetic diversity is of fundamental importance for breeding, as genetic improvement in traits is directly linked to genetic variation (Falconer and Mackay, 1996). Quantitative genetics suggest that both genetic variation and selection efficiency are highest when allele frequencies are intermediate. Selection in closed populations tends to increase inbreeding with a loss of genetic diversity, thereby limiting the long-term response to selection. However, at some point, it is possible to create genetic variability by crossbreeding lines from different genetic backgrounds. To create a new line, the chosen breeds must have complementary traits (Legault et al., 1996). In general, the new line is closed after the original crossbreeding takes place, meaning reproducers are only chosen among crossbred animals. After several generations, the population becomes genetically homogeneous and can be considered as a new line. One of the expected results of this process is an increase in the genetic diversity of the crossbred animals (Bidanel, 1992). Analyses of French and Spanish pure and composite pig lines using microsatellite markers showed high within-population variability in composite lines compared to pure lines Boitard et al. (2010). In addition, composite lines were sufficiently different from the pure breeds to be identified as separate populations.

The first generations of a synthetic line are usually managed without selection in order to promote the mixing of genomes from pure breed parental populations and to preserve the original genetic diversity. However, few data are available regarding how the genetic diversity of the first generations of an animal composite line is managed. The lack of data is due to uninformative pedigree data because the genealogical relationships among founder breeds used in the crossbreeding were not established, making it impossible to characterize the genetic diversity in crossbred animals. To overcome this problem, genomic data could be analyzed to characterize the resulting genetic diversity (Zhang et al., 2019) and new strategies could be designed to characterize and promote genetic diversity right at the beginning of a new synthetic line (Gobena et al., 2018; McTavish and Hillis, 2014).

A specific breeding scheme can be established after two or three generations of mixing (Legault et al., 1996). However, the original genetic diversity has to be preserved in order to promote the complementarity of the genomes of the original breeds and to limit genetic drift towards one of the parent breeds, particularly through selection (Paim et al., 2020). Knowledge of the genomic composition of crossbred animals would thus help keep the genomic composition between the parental breeds balanced over generations. The mating plan for breeding stock is also an important step to favor haplotype recombinations and limit inbreeding. Inbreeding can be characterized at the genomic level by different methods. One of them is the detection of runs of homozygosity (ROH), continuous homozygous chromosomal segments along the genome (Peripolli et al., 2017). An interesting approach was proposed by de Cara et al. (2013) to compute a coancestry coefficient based on ROH named *f*_*seg*_. In each male/female pair, the coancestry coefficient reflects the expected generation of ROH in any potential offspring. Choosing animal pairs with low *f*_*seg*_ should limit ROH-based inbreeding in the next generation.

The two main objectives of the present study were first, to assess a strategy for the management of genetic diversity in the first generations of a composite line, and second, to evaluate the impact of the number of generations before the start of selection on the ongoing genetic diversity and genetic progress.

## 2 Material and methods

Stochastic simulations were used to compare various breeding strategies using the R-package MoBPS (Pook et al., 2020). The code exemplary needed to perform the simulations is provided in **Supplementary File S1**.

### 2.1 Genomic and phenotypic data

For the simulations, the base pig population was created using real genomic and phenotypic data collected from a three-way crossbreeding program previously described in Ganteil et al. (2021). In the first generation, Large White sows were mated with Pietrain boars. In the next generation, Pietrain x Large White sows (PLW) were mated with Duroc boars. The offspring of the second crossbreeding are named G0.

Genomic data were obtained from the breeding company NUCLEUS. The reproducers from the G0 population, 82 boars and 676 sows, were genotyped with the Illumina Porcine Chip, Porc XT 60K. The reference map was based on the *Sus scrofa* 11.1 pig genome assembly. Quality control of genotypes was performed with PLINK v1.9 software (Chang et al., 2015). Only markers on autosomes were kept. Markers with more than 5% of missing genotypes were discarded. Minor allele frequency (MAF) pruning was performed to eliminate alleles with a frequency below 0.01%. After quality control, 758 animals and 46 628 SNP were used for analysis. Genotypes were then phased with the `BEAGLE` v5.1 software (Browning et al., 2018).

Phenotypic data on the G0 population were collected by the breeding company NUCLEUS. To simplify the simulations, only one trait was considered for selection: the age of the pigs at 100 kg (A100), an indicator of growth rate. The animals were weighed at around 150 days old, after which A100 was computed using the following formula:

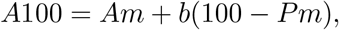

where *Am* is the age of the animal at weighing and *Pm* the measured weight. Coefficient *b* was estimated by IFIP-*Institut du porc* (Jourdain et al., 1989). For these animals, *b* was calculated using the following formula: *b* = *C* − 0.0077 × *Wb* + 0.0047 × *Ab*, where *C* = 1.075, *Wb* is the mean weight of the batch and *Ab* is the mean age of the batch. To generate a simulated trait architecture that is realistic, all the SNP included were assigned linear effects based on the estimation computed by ridge regression BLUP (Akdemir and Godfrey, 2015). This method is available with `MoBPS`.

### 2.2 Population structure in simulations

All the simulation scenarios are presented in **Figure 1**. In the first step, we simulated three different scenarios to compare three breeding strategies. In these scenarios, the starting population was G0 with 10 subsequently simulated generations. The size of the population and selected reproducers were chosen to fit real pig populations. At each generation, 30 males and 300 females were selected as reproducers. Each male was mated with 10 females. The mating plan differed between the scenarios, as it will be explained in the next subsection. Multiple offspring were produced by the same sow/boar pair to obtain 1 000 males and 1 000 females. To account for the variability due to random chromosomal segregation and recombinations during meiosis, scenarios were simulated either 10 or 25 times. The number of repetitions has been defined according to our computation capacities.

**Figure 1:**
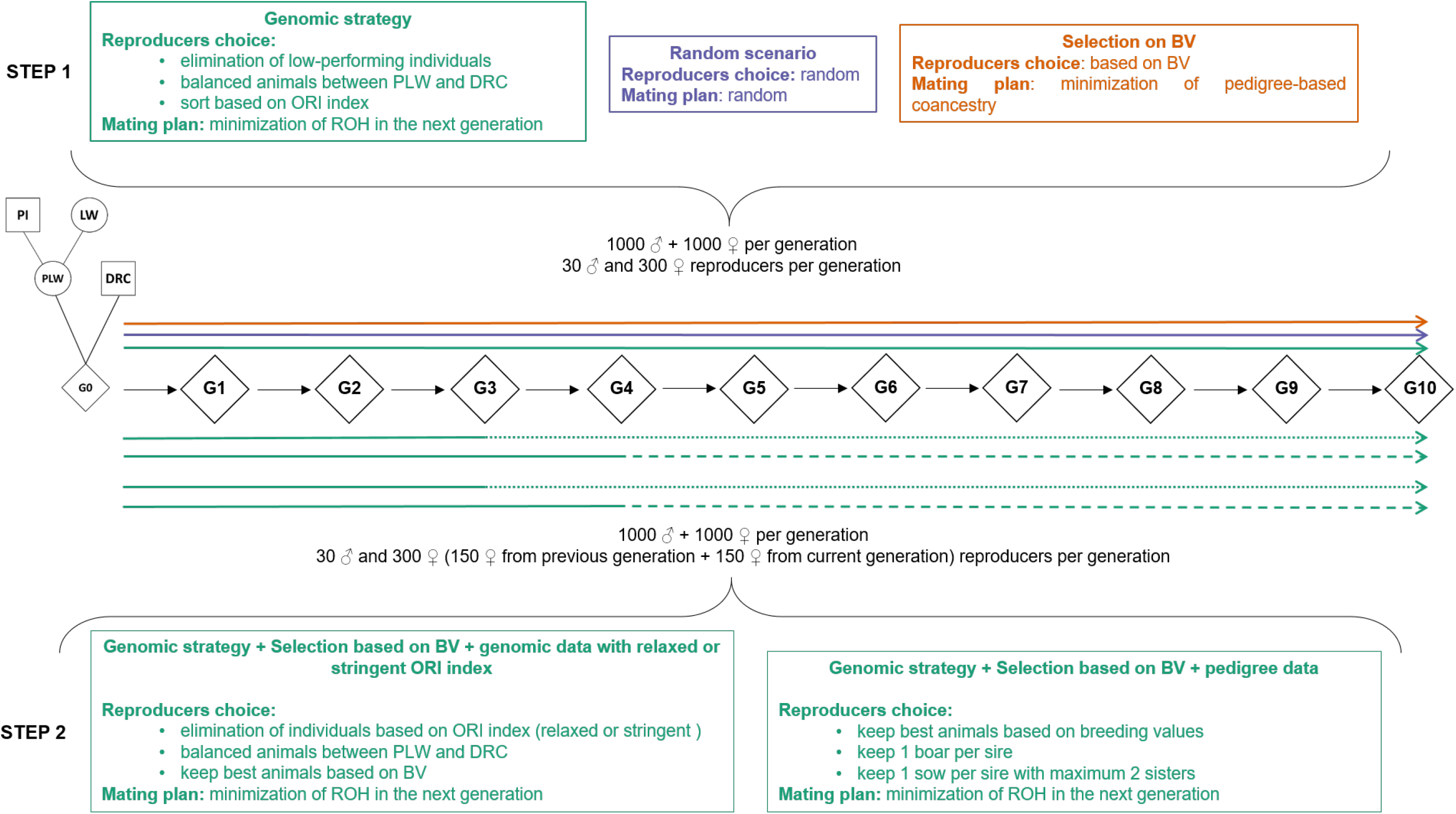
Simulation scenarios. In step 1 we compared our diversity management strategy hereafter named ‘Genomic strategy’, with a ‘Random scenario’ and a ‘Selection on BV’ scenario. In step 2, we tested two selection scenarios with two selection starting points, i.e., G3 and G4. (BV: breeding values, DRC: Duroc, PLW : Pietrain x Large White, ORI index: originality index)

In the second step, we compared two different starting times for selection and three selection strategies. The generations before selection started were simulated with the genomic diversity management strategy. We tested two starting generations, G3 and G4, and generated respectively 7 and 6 additional generations. Population size was the same as mentioned above with overlapping generations for females. At each generation, we selected 30 males from the current generation and 150 females from both the previous and the current generation as reproducers. This method was applied from the second selected generation in the simulations.

### 2.3 Evaluation of a genomic strategy to manage genetic diversity

Three different scenarios were computed to compare a strategy using genomic data to maximize diversity with two alternative scenarios.

#### 2.3.1 Random scenario

The first scenario is a basic simulation with both reproducers and mating selected randomly, called **Random scenario**. We repeated this scenario 25 times.

#### 2.3.2 Scenario with selection based on breeding values

In the second alternative scenario, called **Selection on BV**, breeding values (BV) were estimated with `MoBPS` from marker effects estimated for the G0 population. BV were estimated based on the custom implementation of gBLUP (Meuwissen 2001) in `MoBPS` with known heritabilities. We selected reproducers with the lowest BV because the objective of selection was to improve the growth rate by minimizing A100. We computed a coancestry coefficient *f*_*PED*_ based on pedigree data with function `pedIBD()` in the `optiSel` R package (Wellmann, 2019). This coefficient was based on the pedigree of real G0 animals (over 10 generations) concatenated with the simulations without errors. A mating plan was established with an optimization algorithm (in-house `Fortran` script) based on the minimization of *f*_*PED*_ sum between matings. This scenario represented a selection strategy that maximizes genetic gain, where genomic data were used to obtain accurate BV, and mating was managed by minimizing pedigree relationship. **Selection on BV** was repeated 25 times.

#### 2.3.3 Scenario with genomic data to manage genetic diversity

The third scenario, called **Genomic strategy**, is based on a strategy where genomic data are used both for selection and genetic diversity management. Two principles were combined to select reproducers. The first was to choose reproducers among animals with balanced genomic composition between founding breeds, with a particular focus on the proportion of Duroc origin. The second principle was to promote original animals in terms of alleles. An animal is original if it carries alleles that are uncommon in the population.

Genomic compositions between PLW and Duroc were calculated based on the real points of recombination in each meiosis. Note that this is not possible in practice and needs to be estimated and thus represents an idealized estimate compared to a real-world scenario. If a chromosomal fragment originated from the maternal chromosome of a G0 animal, its origin was a PLW founder. Otherwise, if a chromosomal fragment originated from a paternal chromosome of a G0 animal, its origin was a Duroc founder. We considered an animal to be balanced when the proportions of PLW and Duroc origins were between 0.30 and 0.70. For original animals, Danchin-Burge et al. (2016) proposed the computation of an originality index (ORI) to highlight individuals with uncommon alleles compared to a target population. An original animal has a positive ORI index, whereas an animal with common alleles has a negative ORI index. For each animal *i*, ORI index was computed with in-house `Python` script with the following formula:

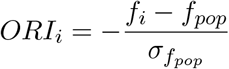

where *f*_*i*_ is the mean allele frequency of markers carried by individual *i, f*_*pop*_ is the mean frequency of all individuals in the population and *σf*_*pop*_ is the standard deviation.

Last, BV were estimated as described in the subsection 2.3.2. Reproducers were selected in three consecutive steps. First, individuals with a very low growth rate, i.e., with BV greater than three standard deviations from the mean, were eliminated. Second, only animals with balanced PLW and Duroc genomic origins were kept. Third, among the remaining animals, the 30 males and 300 females with the highest ORI indexes were selected as reproducers.

The mating strategy was based on the limitation of ROH generation in the offspring. Shared genomic segments between each male/female pair were detected with `GERMLINE` software (Gusev et al., 2009). We used the following parameters: minimum size of a shared segment, 1 Mb and a minimum of 30 SNP per segment. We did not allow mismatching homozygous or heterozygous markers for a shared segment. The ROH coancestry coefficient *f*_*seg*_ (de Cara et al., 2013), which reflects the potential of generation of ROH in the offspring for each pair of individuals was computed as:

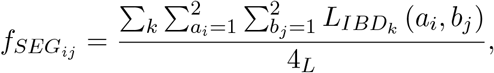

where *L*_*IBD*_ (*a*_*i*_, *b*_*j*_) is the length of the *k*^*th*^ shared segment measured over homologous *a* of animal *i* and homologous *b* of animal *j. L* is the total length of the autosomes covered by markers in bp.

Like in the **Selection on BV** scenario, the mating plan was established with an optimization algorithm. In this scenario, the mating plan was based on the minimization of *f*_*seg*_ sum between mates. This scenario was repeated 10 times.

### 2.4 Comparison of two starting generations for selection

We tested two different selection scenarios with two different starting points. We took the simulated data from the previous **Genomic strategy** scenario, and applied selection criteria with a starting point at G3 and G4. In all scenarios, BV were estimated as previously described in subsection 2.3.2. Each scenario was repeated 10 times.

#### 2.4.1 Selection based on breeding values and pedigree data

This scenario, called **BV + pedigree data** represented a strategy where genomic data are used to accurately estimate BV but not manage diversity. For the choice of reproducers, animals with a high growth rate according to BV were selected with simple constraints based on family structure: one male per sire, a minimum of one female per sire and a maximum of two full sisters. For mating, we assumed that reproducers were genotyped and we mated reproducers as described in the subsection 2.3.3, with minimization of *f*_*seg*_ sum.

#### 2.4.2 Selection based on breeding values and genomic data with a relaxed or stringent ORI index

We simulated a scenario which included two sub-scenarios: **BV + genomic data with relaxed ORI** and **BV + genomic data with stringent ORI**. The difference between these two sub-scenarios was the constraint on the ORI index. For these two sub-scenarios, we applied a step-by-step selection to select reproducers. First, we sorted the animals based on the ORI index. In **BV + genomic data with relaxed ORI**, we kept animals with ORI index greater than -1.5. In **BV + genomic data with stringent ORI**, we kept 25% of males and 66% of females with the highest ORI index. The following steps in reproducer selection were the same in the two sub-scenarios. We kept only animals balanced PLW and Duroc genomic origins and then selected animals with the best BV. For mating, we used genomic data to design a mating plan based on minimization of *f*_*seg*_ sum as described above in **Genomic strategy**.

### 2.5 Data analysis

#### 2.5.1 Allele frequency

Allele frequencies were computed with `PLINK` software. According to their MAF, markers were divided into four categories: Fixed if *MAF* = 0, Rare if *MAF* < 0.05, Intermediate if 0.05 < *MAF* < 0.10 and Common if *MAF >* 0.10. We computed the proportion of alleles in each category.

#### 2.5.2 ROH-based inbreeding

ROH was detected with `PLINK` software based on the same parameters defined in Ganteil et al. (2021). The values selected to define a ROH were 30 SNP and 1,000 kb and the minimum density was set at one SNP per 100 kb. One missing SNP was allowed per sliding window. To obtain strictly homozygous ROH, no heterozygous SNP were allowed per sliding window. All the other parameters available in `PLINK` that are not mentioned above were default settings. We then computed a ROH-based inbreeding coefficient for each animal, *F*_*ROH*_ with `detectRUNS R` package (Biscarini et al., 2019) as:

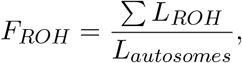

where *L*_*ROH*_ is the sum of the length of all the ROH detected in an animal in bp, and *L*_*autosomes*_ is the total length of the autosomes covered by markers in bp.

#### 2.5.3 Analyses based on identity by state matrix

`PLINK`, was used to compute an identity by state (IBS) matrix to obtain molecular coancestry coefficients *f*_*Mij*_ between each pair of individuals *i* and *j*. The average molecular pairwise coancestry *f*_*M*_ was used to compute expected heterozygosity (*He*) as *He* = 1 *− f*_*M*_ . This parameter can be considered as an indicator of the ability of a population to respond to selection in the short term. Molecular inbreeding coefficients *F*_*Mi*_ (proportion of homozygous markers) were also obtained with `PLINK` for each individual *i*.

#### 2.5.4 Genetic diversity analysis with pedigree data

This analysis was based on the pedigree of real G0 animals concatenated with the pedigree extracted from `MoBPS` for simulated animals. We calculated with `pedInbreeding()` function from `optiSel R` package pedigree-based inbreeding coefficients *F*_*PEDi*_ for each individual *i*.

#### 2.5.5 Breeding values analysis

For each simulated animal, a BV was generated by `MoBPS`. We collected all BV to monitor their evolution and calculated the mean and standard deviation per generation for all 2,000 simulated animals and for all repetitions.

## 3 Results

### 3.1 Genetic diversity management

We compared a diversity management strategy, **Genomic strategy** with two other scenarios **Random scenario** and **Selection on BV** over 10 generations.

#### 3.1.1 Allele frequencies

First, we observed the evolution of allele frequencies (**Figure 2**). For the Fixed category (*MAF* = 0), in all scenarios, the G0 generation did not have fixed alleles because of the pruning previously applied to genomic data. As expected, in the three scenarios, the main class of alleles was the Common class (*MAF* > 0.10). The proportion of this category, decreased more rapidly in the **Selection on BV** scenario than in the two other scenarios. For the other frequency classes, the **Selection on BV** scenario showed a bigger increase in the proportions of Intermediate (0.05 < *MAF* < 0.10), Rare (*MAF* < 0.05) and Fixed classes than in the two other scenarios. These results revealed a greater and faster loss of genetic diversity in the scenario with selection. The results of the **Genomic strategy** and **Random scenario** were close but the decrease in proportion of Common class was greater in the **Random scenario** than in the **Genomic strategy** scenario.

**Figure 2:**
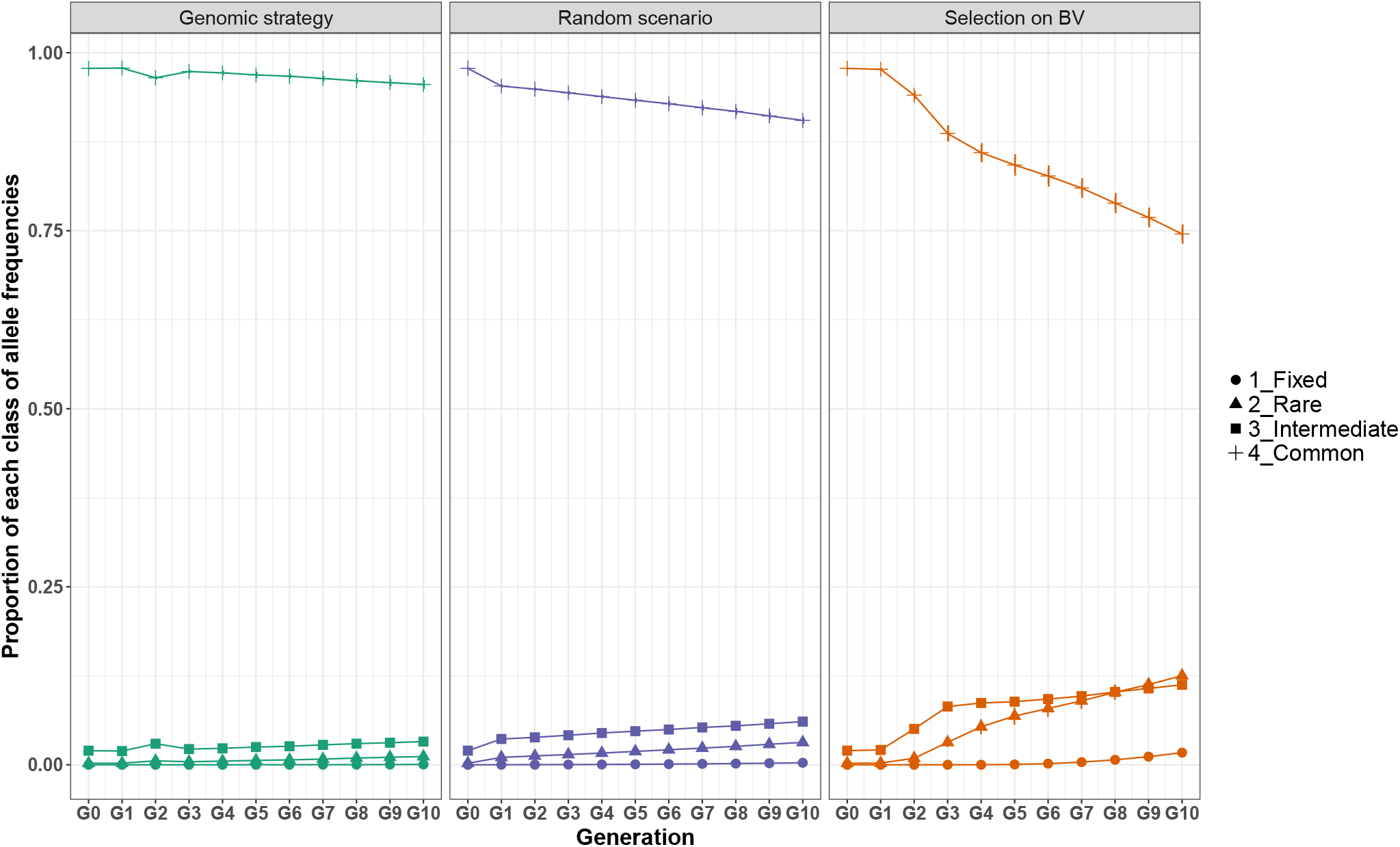
Changes in proportion of each class of allele frequencies over 10 generations in the **Genomic strategy, Random scenario** and **Selection on BV**. Bars indicate the standard deviation across replicates. (Fixed: *MAF* = 0, Rare: *MAF* < 0.05, Intermediate: 0.05 < *MAF* < 0.10 and Common: *MAF >* 0.10, BV: breeding values)

#### 3.1.2 Inbreeding

We analyzed the evolution of inbreeding with ROH-based inbreeding coefficient *F*_*ROH*_, IBS-based inbreeding coefficient *F*_*M*_ and a pedigree-based inbreeding coefficient *f*_*PED*_ in the **Genomic strategy** scenario and in the two alternative scenarios (**Figure 3**).

**Figure 3:**
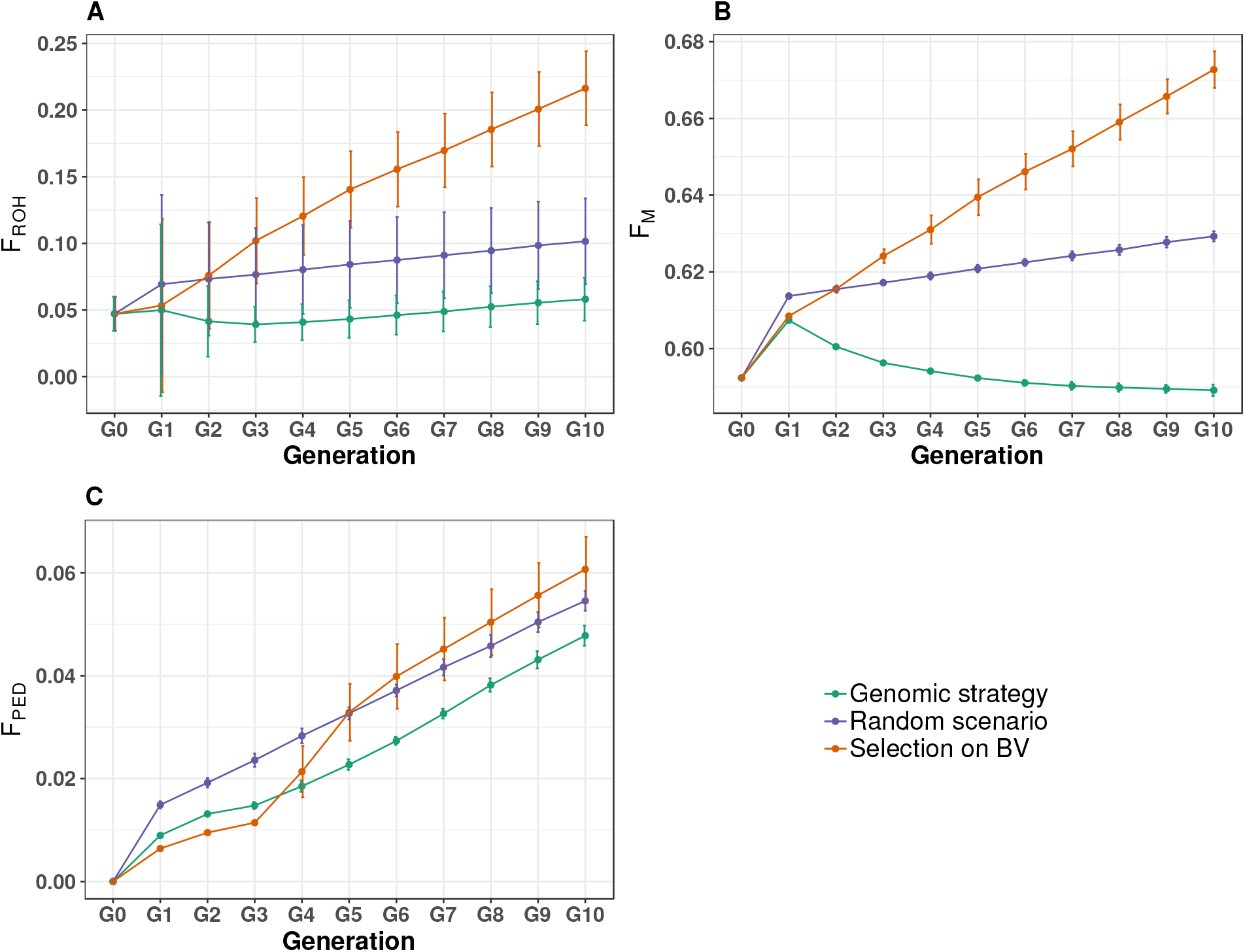
Changes in three inbreeding coefficients, *F*_*ROH*_ (**A**), *F*_*M*_ (**B**), *F*_*PED*_ (**C**) over 10 generations in the **Genomic strategy, Random scenario** and **Selection on BV** scenarios. Bars indicate the standard deviation across replicates. (BV: breeding values)

The two graphs with genomic coefficients *F*_*ROH*_ and *F*_*M*_ were similar (**Figure 3A,B**). They showed a significant increase in the two coefficients in the **Selection on BV** scenario. Inbreeding increased in the **Random scenario** but to a lesser extent than in the **Selection on BV** scenario and its increase was strictly linear from G1 to G10. The most limited increase in inbreeding was observed in the **Genomic strategy** scenario. *F*_*ROH*_ increased slightly between G0 and G1 and decreased slightly after G1 and then increased from G3 to G10. *F*_*M*_ increased between G0 and G1 like *F*_*ROH*_ but after G1, it decreased until G7 and then appeared to stabilize. In G10, *F*_*ROH*_ was 0.21, 0.10 and 0.06 for **Selection on BV, Random scenario** and **Genomic strategy** scenarios, respectively. *F*_*M*_ was 0.67, 0.63 and 0.59 in the **Selection on BV, Random scenario** and **Genomic strategy** scenarios, respectively.

Unlike *F*_*ROH*_ or *F*_*M*_, *F*_*PED*_ increased significantly in all three scenarios **Figure 3C**, albeit with a slightly different pattern. In the **Selection on BV** scenario: between G0 and G3, *F*_*PED*_ remained the lowest, with a bigger increase after G3 to reach the highest *F*_*PED*_ value (0.06) in G10. In the **Random scenario**, *F*_*PED*_ presented a strictly linear increase after G1. The profile of the **Genomic strategy** was similar to that in the **Selection on BV** between G0 and G3, subsequently, *F*_*PED*_ increased at a slower rate and at G10, this scenario had the lowest *F*_*PED*_ value. Standard deviations in *F*_*PED*_ were the highest in **Selection on BV**, especially between G4 and G10.

#### 3.1.3 Expected heterozygosity

In all three scenarios, we observed an increase in *He* at G1 that was higher in the **Genomic strategy** and **Selection on BV** scenarios (**Figure 4**). In the following generations, *He* decreased in **Random scenario** and **Selection on BV**. In **Random scenario**, the decrease was linear and lower than in **Selection on BV** which presented the largest decrease. For **Genomic strategy**, after G1, EH decreased only during 1 generation and increased after G3 and it seemed to stabilize from G3 to G10 around 0.31. We observed standard deviations in *He* in the three scenarios with the highest value in **Selection on BV**.

**Figure 4:**
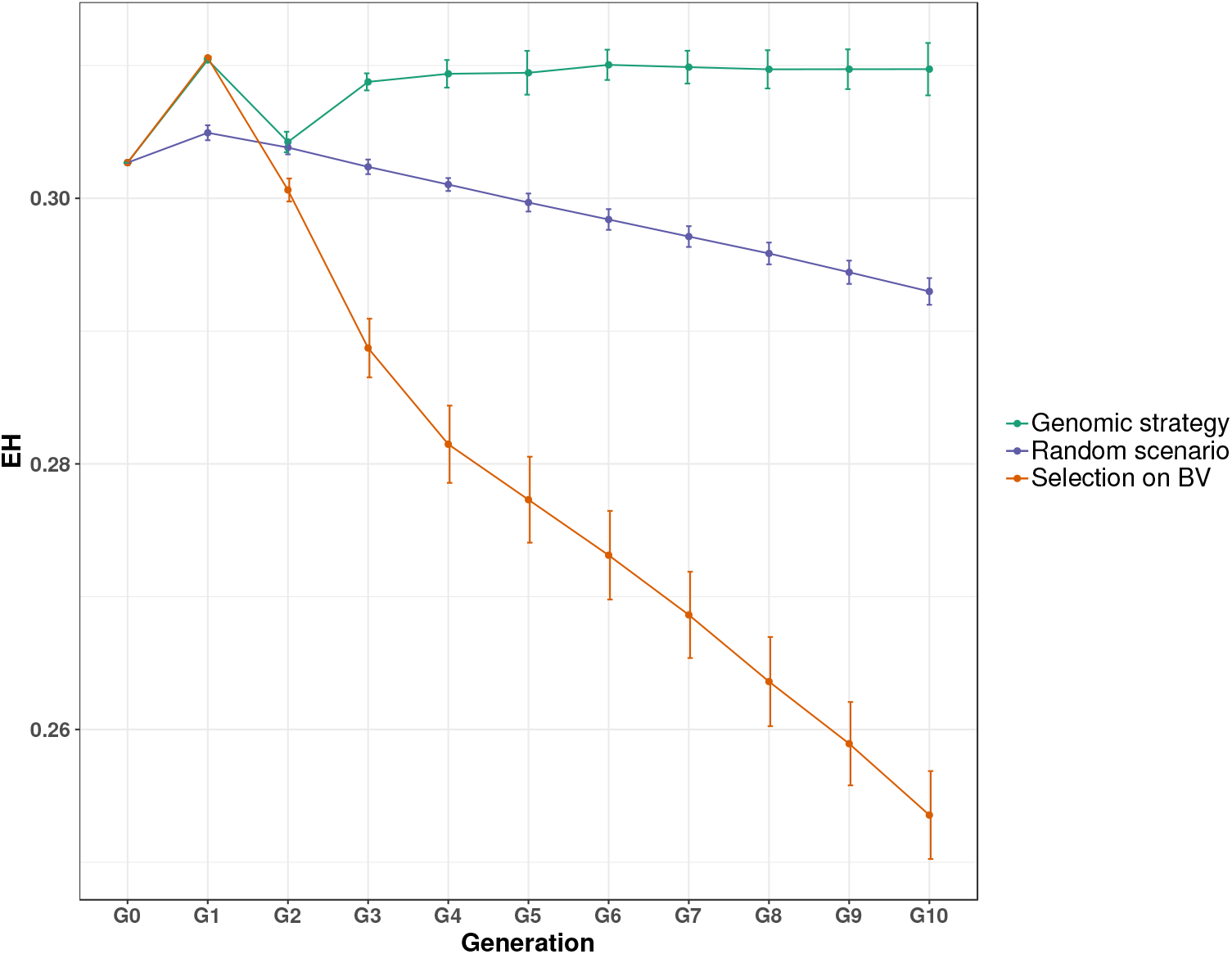
Changes in expected heterozygosity (He) over 10 generations in the **Genomic strategy, Random scenario** and **Selection on BV** scenarios. Bars indicate the standard deviation across replicates. (BV: breeding values)

### 3.2 Impact of starting selection in G3 or G4 on diversity and genetic progress

Here we compared three selection scenarios using two different generations to start the selection, G3 and G4. Genetic diversity estimators and BV were analyzed from G3 to G10 or G4 to G10, respectively.

#### 3.2.1 Allele frequencies

**Figure 5** shows the proportion of Common, Intermediate, Rare and Fixed alleles at each generation. In **BV + genetic data with relaxed ORI**, the proportion of Common alleles decreased and the proportion of Intermediate and Rare alleles increased. In **BV + genetic data with stringent ORI**, the proportion in each allele frequency classes remained relatively constant over generations, underlining the ability of the strategy to conserve genetic diversity. Differences between starting selection in G3 or G4 were negligible in the two scenarios. The shape of the two respective curves for the allele frequency contributions was similar at both starting points, with the G3 curve basically shifted by one generation (to the left) relative to the G4 curves.

**Figure 5:**
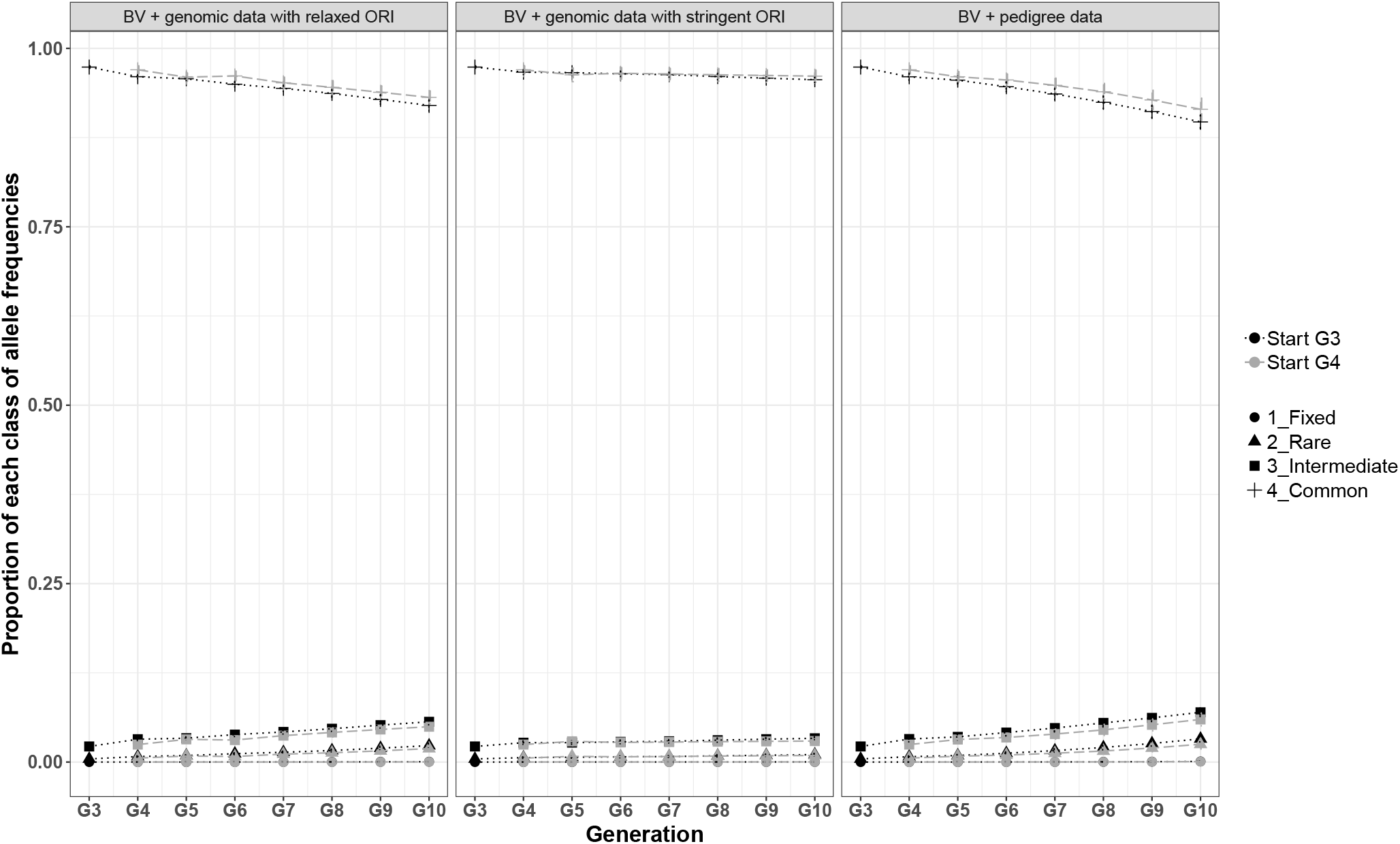
Changes in the of proportion of each class of allele frequencies for **BV + genomic data with relaxed ORI, BV + genomic data with stringent ORI** and **BV + pedigree data** and comparison of selection starting at G3 and G4. Bars indicate the standard deviation across replicates. (Fixed: *MAF* = 0, Rare: *MAF* < 0.05, Intermediate: 0.05 < *MAF* < 0.10, Common: *MAF >* 0.10 and BV: breeding values)

The third scenario **BV and pedigree data**, showed the strongest decrease in the proportion of Common alleles compared to the two other scenarios with genomic data and ORI index. The curves for the start of selection at G3 and G4 were again similar, however, as the shift per generation in **BV and pedigree data** was highest the biggest differences in G10 were observed. There was a slight increase in Intermediate and Rare alleles that was similar in the two starting points.

The proportion of Fixed alleles, was constant in the three scenarios.

#### 3.2.2 Inbreeding

The evolution of inbreeding was characterized with the simulation of selection scenarios. As mentioned above, we studied *F*_*ROH*_, *F*_*M*_ and *F*_*PED*_ (**Figure 6**). **Figure 6A** shows the results for *F*_*ROH*_ were similar in the two scenarios with genomic data and ORI index. *F*_*ROH*_ first increased, then decreased, and then increased again. The shape of the curve was the same whether selection started at G3 or G4, the two curves only shifted by one generation. However, at G10, in the two scenarios, the simulation that started at G3 had a higher *F*_*ROH*_ than the simulation that started at G4. Regarding the difference between the scenarios **BV + genomic data with relaxed ORI** and **BV + genomic data with stringent ORI**, in limiting inbreeding, the second scenario was more efficient. In the third selection scenario, **BV + pedigree data**, *F*_*ROH*_ increased immediately after the start of selection until G10. The increase was lower when selection started in G4 than when it started in G3.

**Figure 6:**
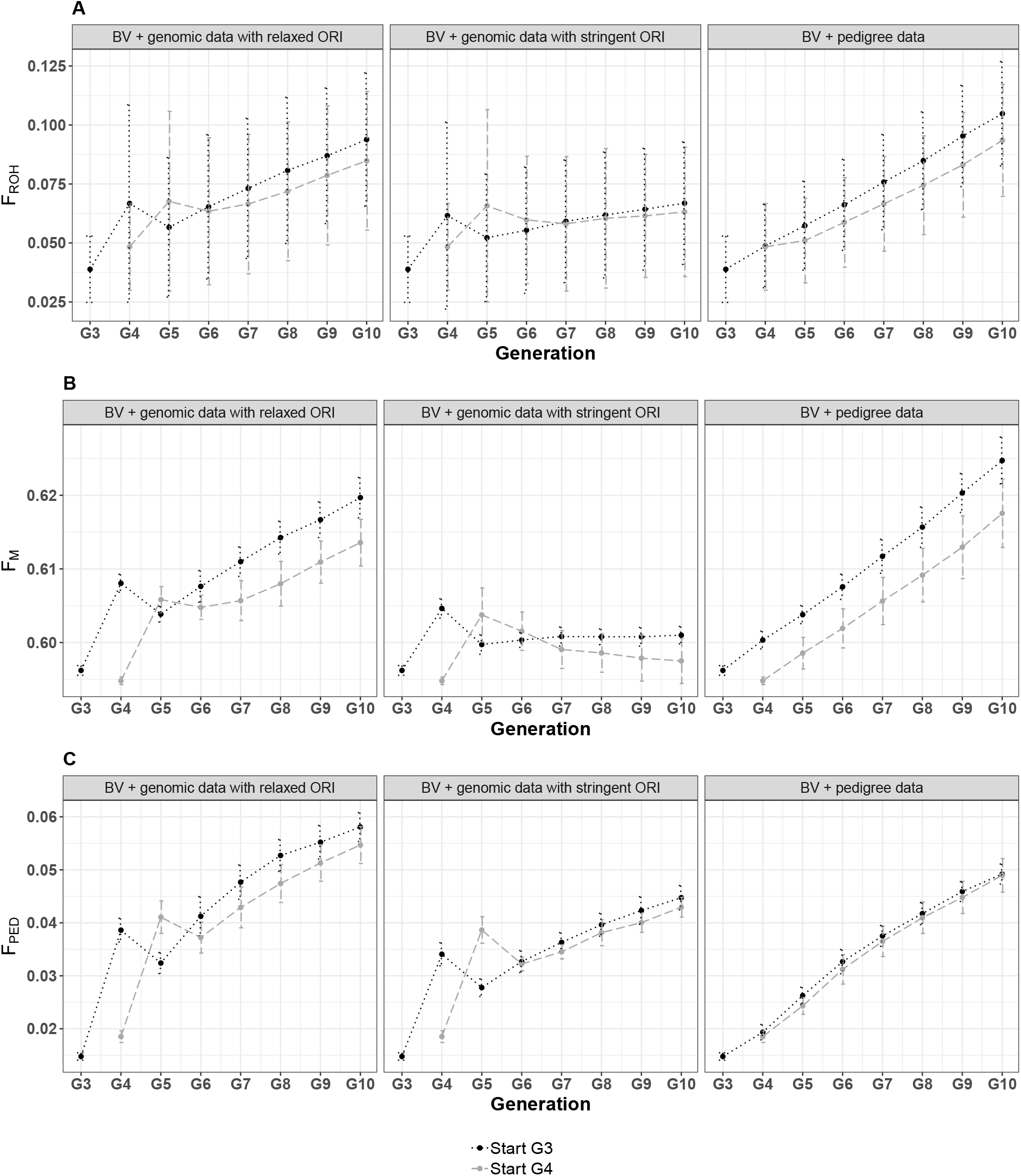
Evolution of inbreeding in **BV + genomic data with relaxed ORI, BV + genomic data with stringent ORI** and **BV + pedigree data** and comparison of selection starting at G3 and G4. The three inbreeding coefficients analyzed were *F*_*ROH*_ (**A**), *F*_*M*_ (**B**) and *F*_*PED*_ (**C**). Bars indicate the standard deviation across replicates. (BV: breeding values)

The second genomic-based inbreeding coefficient was *F*_*M*_ (**Figure 6B**). The results for this coefficient were similar to previous results for *F*_*ROH*_. Once again, starting selection at G4 limited the increase in inbreeding better.

Last, we studied *F*_*PED*_ (**Figure 6C**), the shapes of the curves in the scenarios with genomic data with ORI index were similar. The results of the scenario **BV + pedigree** were very close results for the two selection starting points.

For the three coefficients, the scenario **BV + genomic data with stringent ORI** limited inbreeding the most. However, in the first generations of selection (G4 to G6), the scenario **BV + pedigree** with a start of selection at G4 showed the lowest level of inbreeding.

#### 3.2.3 Evolution of breeding values

The estimated genetic progress was compared in the three selection scenarios (**Figure 7**). The selection applied in the three scenarios aimed to decrease the BV of the studied trait (A100). In all scenarios and in line with the breeding objective, BV decreased over the 6 or 7 simulated generations. These results highlighted improved growth of simulated animals. In the **BV + genomic data with relaxed ORI** and **BV + pedigree data** scenarios, there was a shift of approximately one generation between starting selection at G3 and at G4 in BV evolution. This result was not observed for **BV + genomic data with stringent ORI** where the improvement of BV from G7 was somewhat limited when selection started at G4.

**Figure 7:**
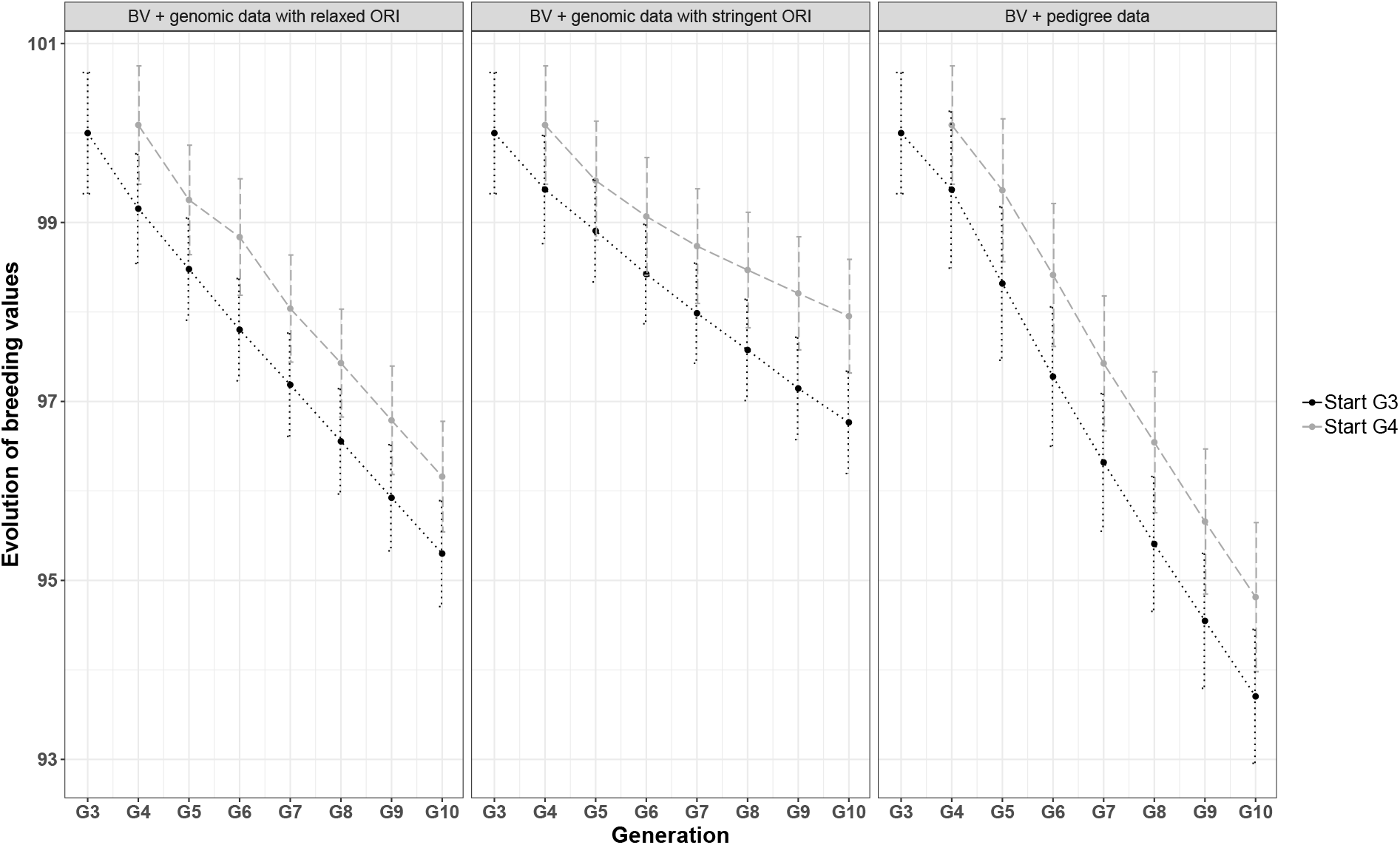
Evolution of BV for **BV + genomic data with relaxed ORI, BV + genomic data with stringent ORI** and **BV + pedigree data** with comparison of selection starting at G3 and G4. The trait studied here was age at 100 kg (A100) to measure the efficiency of growth. (BV: breeding values)

As expected, in G10, the best improvement of BV was observed for **BV + pedigree data**, where in simulations when selection started at G3, the mean of the BV was 93.8 compared with 95.4 and 96.8 in the **BV + genomic data with relaxed ORI** and **BV + genomic data with stringent ORI** scenarios, respectively.

## 4 Discussion

### 4.1 Simulations using real data

This article describes a simulation study to compare different strategies to create a synthetic line in pig breeding. Founder animals are generated based on real genomic data and the trait is generated using real phenotypic data. The G0 population is the product of two steps three-way crossbreeding and G0 animals constitute the first generation of a new synthetic line. We started all simulations using all G0 reproducers’ genotypes and phenotypes available. Additional genomic data for G1 and G2 animals were also used and we were thus able to compute *F*_*ROH*_ (**Supplementary File S2**). To validate our simulation approach, changes in *F*_*ROH*_ were compared between the real population and the simulated data in the scenario **Genomic strategy** for the first three generations. This scenario was based on the strategy carried out on the line management. We found high similarity between the evolution of real and simulated inbreeding coefficients, thus confirming the relevance of our simulation analyses.

Population characteristics such as the number of offspring and the number of selected reproducers or management system (like overlapping generations) were chosen based on the French porcine industry. We tested different simulation scenarios to manage diversity, with and without genomic data. Our aim was to explore the possibility of diversity management offered by genomic data. We consider scenarios with no genomic data or only designed to manage matings by breeders, close to current practices as genotyping of all candidates for selection is usually considered too costly.

In the pig breeding literature, simulation studies are generally based on fully simulated data. For instance, Gourdine et al. (2012) simulated all data of a small local pig breed to explore genetic diversity management with optimal contribution selection (OCS). When all data are simulated, it is possible to test many simulation scenarios with different parameters for example population size or selection pressure. Simulations based on real data should be more useful to monitor the evolution of a specific population. Bosse et al. (2015) compared different strategies in two different populations, a local breed *Sus cebifrons* and commercial breed Pietrain. Bosse et al. (2015) studied real genomic data and simulated the management of each population. They confirmed previous results obtained with simulated data by de Cara et al. (2013) about the efficiency of *f*_*seg*_ to mate animals.

However, the simulated scenarios have some limitations. First, to explore genetic progress, BV were computed for only one trait, A100, which estimated growth efficiency using real phenotypic data measured in G0 animals. So, were close to a real trait but in practice, selection is done used a selection index and/or based on multiple traits. In that respect, the simulation scenarios are a simplistic representation of the breeding objective of a sire line. Moreover, we did not account for potential differences in genetic levels between the three founder populations. With only simulated data, we could have simulated different genetic backgrounds for the founder populations but we had no data available to calibrate such differences in a realistic way. However, differences between using an index or a trait should be limited and the main conclusion of our study should still hold valid. In theory, simulation of these factors in `MoBPS` would be possible.

Another limitation of our study is the contemporaneity of all the candidates. We assumed that all candidates for selection were available at the same time and could thus be chosen as reproducers and mated simultaneously. However, this is not the case in breeding pig populations, some matings cannot be conducted due to the spacing out of the animals from one generation.

### 4.2 Maximization of genetic diversity in the first generations of a new line

Maximizing genetic diversity is an important step during the first generations of a new line before starting selection but few data are available on this topic in the context of the creation of a new line. The increase in genetic diversity in the composite line compared to the founding breeds is expected from the increase of heterozygosity. With pedigree relationships only, inbreeding is assumed to be nonexistent in the first crossbreeding generation, which can lead to overestimation of genetic diversity. Using genomic data, it is possible to characterize genetic diversity with better accuracy. Some studies have shown genomic inbreeding could persist in crossbred animals (Ganteil et al., 2021; Howard et al., 2016).

For this reason, we proposed the **Genomic strategy** as an innovative way to manage genetic diversity in the first generations of a new pig line. Our scenario included genomic data for the choice of reproducers and their mating. We estimated two genomic criteria to select reproducers: the ORI index and balanced animals for one of the three founding populations. The ORI index makes it possible to enhance animals with uncommon alleles in each generation and helps maintain rare alleles in the population. In the long term, using the ORI index, we expect to obtain animals with ORI index close to 0 which means an increase in heterozygosity in animals. The second criterion based on the origin of the founders’ genomes in crossbred animals limits genetic drift towards a particular founder breed. As we did not introduce different genetic levels between the founder breeds for the selected trait, no such systematic breed favoritism was expected, except by chance. Using these genomic criteria to select reproducers combined with minimization of *f*_*seg*_ coefficient in mating plans are efficient ways to maintain genetic diversity compared to a random choice of reproducers and their matings, like in the **Random scenario**.

In our simulations, the choice of balanced animals was only based on the estimation of the Duroc and PLW origins. The method could be improved by estimating all the origins (Duroc, Large White and Pietrain). For this purpose, a robust method could be to phase each genotype of the crossed animals and identify the origin of each chromosomal fragment down to the purebred founders to assign each fragment to a Pietrain, Large White or Duroc origin.

To mate animals, we chose a method based on the minimization of *f*_*seg*_ coefficient which limits ROH in the offspring (de Cara et al., 2013). de Cara et al. (2013) performed simulation analyses and suggested that their method leads to higher levels of population fitness. This result could be due to the limitation of large ROH by *f*_*seg*_. In fact, large ROH correspond to recent inbreeding (Curik et al., 2014) which is assumed to be more harmful than ancient inbreeding, because selection has had time to reduce the frequency of deleterious alleles that are purged over time (Doekes et al., 2019).

In the **Selection on BV** scenario, the selection of reproducers was only based on BV. In this case, we observed a decrease in the proportion of SNP in Common frequency class, an increase in the proportion of SNP in the three other frequency classes: Intermediate, Rare and Fixed. We chose to analyze allele frequencies by divided them into different frequency classes in order to easily monitor the evolution of frequencies. In the **Selection on BV** scenario, the smallest increase among Intermediate, Rare and Fixed classes, is observed for the Fixed category probably due to MAF pruning applied before starting analysis of the simulation. In our context, i.e, the creation of a new line, the conservation of the specific alleles of each founder breed is important. Our study showed that the **Genomic strategy** could limit the loss of these specific alleles because the evolution of all frequency classes remained almost constant over the 10 generations, in contrast to the **Random scenario** and **Selection on BV** scenario.

When we compared **Selection on BV**, the **Random scenario** and the **Genomic strategy** in terms of inbreeding, the **Genomic strategy** was the most efficient scenario to limit inbreeding. The increase in inbreeding observed in the **Random scenario** was probably due to genetic drift because we analyzed a small population.

Regarding inbreeding estimated with *F*_*PED*_ in the **Selection on BV, Random scenario** and **Selection on BV** scenarios, the results in G10 were similar in the three scenarios, in contrast to *F*_*ROH*_ and *F*_*M*_ . These results confirm the relevance of genomic coefficients to accurately characterize inbreeding by capturing the variation due to Mendelian sampling and thus make it is possible to differentiate animals with the same pedigree (Villanueva et al., 2021). What is moreover, in the context of crossbred animals used in this study, genealogical would not have been informative concerning G0 animals, where all *F*_*PED*_ were equal to 0.

Despite crossbreeding and the generation of heterozygosity, our results show it is not sufficient to prevent an increase in inbreeding when we apply selection like **Selection on BV**. Systems that use maximization of diversity like **Genomic strategy** followed by selection with diversity management using genomic data like **BV + genomic data with relaxed ORI, BV + genomic data with stringent ORI** or a combination of pedigree and genomic like **BV + pedigree data** appear to be appropriate ways to limit an increase in inbreeding. The results we obtained for *He* in the **Genomic strategy** is an additional argument for the use of a diversity management strategy with genomic data during the first generations of a new line. Indeed, in this scenario, *He* was constant after G3. *He* measures the ability of a population to respond to selection in the short term. We hypothesize that starting selection at G3 or at G4 after diversity management, for exampling using the **Genomic strategy** would generate genetic progress.

### 4.3 Comparison of selection strategies

Our objective was to analyze the impact of two alternative selection starting points on genetic diversity and BV evolution. In this context, we studied a breeding program with first generations of the new line managed using a **Genomic strategy**. We then tested a starting selection at G3 and G4 with three different selection scenarios **BV + genomic data with relaxed ORI, BV + genomic data with stringent ORI** and **BV + pedigree data**. It would have been possible to start selection at several generations, but we chose G3 and G4 in accordance with results obtained for *He* which showed stabilization of this indicator after G3. Moreover, in a real synthetic line project, it would be difficult to put off selection to later generations because of breeders’ expectations in terms of genetic progress.

The three selection scenarios we used to compare starting selection at G3 and G4 can be divided into two groups. In the **BV + genomic data with relaxed ORI** and **BV + genomic data with stringent ORI** scenarios, we considered that all the candidates for selection had genomic data whereas in the **BV + pedigree data** scenario, genomic data were available only for selected reproducers. However, in all three scenarios, selection was focused on growth efficiency so minimization of A100’s BV and BV were estimated from genomic data. We made this choice to ensure precise BV and the same method to estimate them in all simulations. So, in the **BV + pedigree data** scenario, the idea was to have genomic data to manage diversity only for reproducers mating.

Regarding diversity parameters like allele frequencies and inbreeding, **BV + genomic data with relaxed ORI** and **BV + genomic data with stringent ORI** proved to be efficient to limit the loss of diversity especially with a stringent selection on the ORI index. The ORI index thus appears to be an easy to use and efficient criterion to preserve diversity in the population. A previous study showed the relevance of the ORI index in cattle compared to using genealogical data (Danchin-Burge et al., 2016).

We observed a particular variation in the inbreeding coefficients after the start of selection in the two scenarios with selection on BV and genomic data with the ORI index. We observed an increase in *F*_*ROH*_, *F*_*M*_ and *F*_*PED*_ between the starting generation of selection and next generation, after which inbreeding decreased and then increased again. In these two scenarios, in the second generation of selection, the choice of female reproducers was conducted with overlapping generations, with half the reproductive females from the previous generation. Complementary analyses were performed in a system with discrete generations and we did not observe this variation at the beginning of selection (data not shown). This phenomenon was not observed in **BV + pedigree data** scenario, so it was probably due to the interaction between ORI index and the start of overlapping generations. In fact, we selected original animals with the ORI index and the selection on original animals was high, but with the selection on BV, we observed an increase in inbreeding. This increase was higher in the **BV + genomic data with relaxed ORI** scenario than in the **scenario BV + genomic data with stringent ORI** scenario because we selected with relaxed ORI, animals with higher BV and maybe genetically closer, the constraint on diversity is less strong than with the stringent ORI selection. The decrease in inbreeding observed after the second generation of selection is probably due to the overlapping of generations. Indeed, overlapping generations promote the minimization of parents’ coancestry and hence reduce inbreeding in offspring (Sonesson and Meuwissen, 2001).

This variation in inbreeding described above was not observed in the scenario **BV + pedigree data**. Using pedigree structure to control for inbreeding was more efficient in limiting inbreeding in the first generations of selection than in the scenarios that use the ORI index. However, after several generations, the two scenarios that use the ORI index were more efficient.

Although **BV + pedigree data** was less efficient in limiting inbreeding than **BV + genomic data with relaxed ORI** and **BV + genomic data with stringent ORI**, all three scenarios respected the recommended rates of inbreeding for small livestock populations. These rates are 0.5-1% rise per generation for *F*_*PED*_ and 0.25-0.50% rise per generation for *F*_*M*_ (Meuwissen and Oldenbroek, 2017).

If we extrapolate these three scenarios to real conditions, scenarios with genomic data available for all candidates for selection have a significant additional cost over the scenario with genomic data available only for reproducers. An alternative could be only genotyping candidates for selection that have interesting phenotypes. In these candidates, diversity criteria like the ORI index or genomic composition in terms of founder origins could be analyzed. These results would help breeders in their final choice of future reproducers.

In the three scenarios in which we tested two selection starting points, our results suggest that, in the context of diversity analysis, it would be advantageous to start selection at G4 rather than G3. One additional generation managed with **Genomic strategy** allowed further chromosomal recombinations that increase diversity within the population. On the other hand, starting selection at G3 is more efficient and generates genetic progress more rapidly than starting at G4. Not surprisingly, the higher the pressure on diversity, the lower the pressure on genetic progress. Therefore, the first three generations of a new line managed with **Genomic strategy** followed by **BV + genomic data with relaxed ORI** appears to be a good compromise between conserving diversity and achieving genetic progress.

We did not analyze a scenario with OCS in this paper. This method of selection uses the average coancestry of the selected parents to manage genetic diversity. This can be implemented in different ways, such as maximizing genetic progress with a fixed rate of inbreeding or minimizing the loss of genetic diversity. OCS can be used with pedigree or genomic data, and is considered as a method of choice for simultaneous improvement of genetic progress and the preservation of diversity (Woolliams et al., 2015). The efficiency of OCS methods is impacted by the estimation of the relationship matrix and it appears that the appropriate genomic relationship matrix should be chosen depending on the diversity targeted (Meuwissen, 2020; Morales-González et al., 2020).

## 5 Conclusion

Genomic data offer new opportunities for genetic diversity management in composite breeds. Our results show that diversity management based on genomic data can be used in the first generations of a new line to build diversity even before starting selection. In the case of two step three-way crossbreeding, it was clearly better to start the selection after 4 generations post-crossbreeding to limit inbreeding, however it is difficult to extend this recommendation to all kinds of crossbreeding strategies. In any case, simulations based on real data collected from the base crossbred population should be carried out to design an appropriate breeding strategy that combines selection and diversity management. Among the breeding scenarios studied here, a strategy that includes a focus on allele frequencies, such as the **BV + genomic data with relaxed ORI** would be a good compromise between genetic progress and diversity conservation.

## Funding

This study was supported by ANRT (Association Nationale Recherche Technologie) with a Doctoral fellowship (2018/0862). This work also received funding from GDivSelGen (Efficient Use of Genetic Diversity in Genomic Selection, Paris, France) action (INRA SelGen metaprogram).

## Acknowledgments

We thank all members of NUCLEUS R&D and technical services as well as the breeders involved in this project. We thank Jean-Michel Elsen for his advice.

## Supplementary Material

- **Supplementary File S1**: Data and scripts can be found at: https://doi.org/10.15454/WWUWDK/M0U6MG
- **Supplementary File S2**: Meeting talk, https://doi.org/10.15454/WWUWDK/NCRDW6

## Notes

https://doi.org/10.15454/WWUWDK

### Competing Interest Statement

The authors have declared no competing interest.

## References

Akdemir, D. and Godfrey, O. U. (2015). EMMREML: Fitting mixed models with known covariance structures.

Bidanel, J.-P. (1992). Comment exploiter la variabilité génétique entre races : du croisement simple àla souche synthétique. INRA Productions Animales, hors série “Eléments de génétique quantitative et application aux populations animales”:249–254.

Biscarini, F., Cozzi, P., Gaspa, G., and Marras, G. (2019). detectRUNS: Detect runs of homozygosity and runs of heterozygosity in diploid genomes.

Boitard, S., Chevalet, C., Mercat, M.-J., Meriaux, J. C., Sanchez, A., Tibau, J., and Sancristobal, M. (2010). Genetic variability, structure and assignment of Spanish and French pig populations based on a large sampling: Variability of Spanish and French pig populations. Animal Genetics, 41(6):608–618.

Bosse, M., Megens, H.-J., Madsen, O., Crooijmans, R. P., Ryder, O. A., Austerlitz, F., Groenen, M. A., and de Cara, M. A. R. (2015). Using genome-wide measures of coancestry to maintain diversity and fitness in endangered and domestic pig populations. Genome Research, 25(7):970–981.

Browning, B. L., Zhou, Y., and Browning, S. R. (2018). A one-penny imputed genome from next-generation reference panels. The American Journal of Human Genetics, 103:338–348.

Chang, C. C., Chow, C. C., Tellier, L. C., Vattikuti, S., Purcell, S. M., and Lee, J. J. (2015). Second-generation PLINK: rising to the challenge of larger and richer datasets. GigaScience, 4(1):7.

Curik, I., Ferenčaković, M., and Sölkner, J. (2014). Inbreeding and runs of homozygosity: a possible solution to an old problem. Livestock Science, 166:26–34.

Danchin-Burge, C., Moureaux, S., Boichard, D., Baur, A., and Fritz, S. (2016). Un indicateur innovant de variabilité génétique àpartir des données moléculaires. In Congrès Mondial de la race Brune, Mende, France.

de Cara, M. R., Villanueva, B., Toro, M., and Fernández, J. (2013). Using genomic tools to maintain diversity and fitness in conservation programmes. Molecular Ecology, 22(24):6091–6099.

Doekes, H. P., Veerkamp, R. F., Bijma, P., de Jong, G., Hiemstra, S. J., and Windig, J. J. (2019). Inbreeding depression due to recent and ancient inbreeding in Dutch Holstein–Friesian dairy cattle. Genetics Selection Evolution, 51(1):54.

Falconer, D. and Mackay, T. (1996). Introduction to quantitative genetics. In Introduction to quantitative genetics. Longman, Harlow England.

Ganteil, A., Rodriguez-Ramilo, S. T., Ligonesche, B., and Larzul, C. (2021). Characterization of autozygosity in pigs in three-way crossbreeding. Frontiers in Genetics, 11:584556.

Gobena, M., Elzo, M. A., and Mateescu, R. G. (2018). Population structure and genomic breed composition in an Angus–Brahman crossbred cattle population. Frontiers in Genetics, 9:90.

Gourdine, J. L., Sørensen, A. C., and Rydhmer, L. (2012). There is room for selection in a small local pig breed when using optimum contribution selection: A simulation study. Journal of Animal Science, 90(1):76–84.

Gusev, A., Lowe, J. K., Stoffel, M., Daly, M. J., Altshuler, D., Breslow, J. L., Friedman, J. M., and Pe’er, I. (2009). Whole population, genome-wide mapping of hidden relatedness. Genome Research, 19(2):318–326.

Howard, J. T., Tiezzi, F., Huang, Y., Gray, K. A., and Maltecca, C. (2016). Characterization and management of long runs of homozygosity in parental nucleus lines and their associated crossbred progeny. Genetics Selection Evolution, 48(1):91.

Jourdain, C., Gueblez, R., and Le Henaff, G. (1989). Ajustement, à poids vif constant, des critères de contrôle en ferme chez le Large-White et le Landrace français. Journées Recherche Porcine, 21:399–404.

Legault, C., Ménissier, F., Mérat, P., Ricordeau, G., and Rouvier, R. (1996). Les lignées originales de l’INRA : historique, développement et impact sur les productions animales. INRAE Productions Animales, 9(HS):41–56.

McTavish, E. J. and Hillis, D. M. (2014). A Genomic approach for distinguishing between recent and ancient admixture as applied to cattle. Journal of Heredity, 105:445–456.

Meuwissen, T. H. and Oldenbroek, K. (2017). Chapter 5. Genetic diversity in small in vivo populations. In Genomic management of animal genetic diversity, pages 139–154. Wegeningen academic publishers edition.

Meuwissen, T. H. E. (2020). Management of genetic diversity in the era of genomics. Frontiers in Genetics, 11:16.

Morales-González, E., Saura, M., Fernández, A., Fernández, J., Pong-Wong, R., Cabaleiro, S., Martínez, P., Martín-García, A., and Villanueva, B. (2020). Evaluating different genomic coancestry matrices for managing genetic variability in turbot. Aquaculture, 520:734985.

Paim, T. d. P., Hay, E. H. A., Wilson, C., Thomas, M. G., Kuehn, L. A., Paiva, S. R., McManus, C., and Blackburn, H. D. (2020). Dynamics of genomic architecture during composite breed development in cattle. Animal Genetics, 51(2):224–234.

Peripolli, E., Munari, D. P., Silva, M. V. G. B., Lima, A. L. F., Irgang, R., and Baldi, F. (2017). Runs of homozygosity: current knowledge and applications in livestock. Animal Genetics, 48(3):255–271.

Pook, T., Schlather, M., and Simianer, H. (2020). MoBPS - Modular breeding program simulator. G3, 10(6):1915–1918.

Sonesson, A. K. and Meuwissen, T. H. E. (2001). Minimization of rate of inbreeding for small populations with overlapping generations. Genetics Research, 77(03).

Villanueva, B., Fernández, A., Saura, M., Caballero, A., Fernández, J., Morales-González, E., Toro, M. A., and Pong-Wong, R. (2021). The value of genomic relationship matrices to estimate levels of inbreeding. Genetics Selection Evolution, 53(1):42.

Wellmann, R. (2019). Optimum contribution selection for animal breeding and conservation: the R package optiSel. BMC Bioinformatics, 20(1):25.

Woolliams, J., Berg, P., Dagnachew, B., and Meuwissen, T. (2015). Genetic contributions and their optimization. Journal of Animal Breeding and Genetics, 132(2):89–99.

Zhang, J., Song, H., Zhang, Q., and Ding, X. (2019). Assessment of relationships between pigs based on pedigree and genomic information. Animal, 14(4):697–705.

